# Cell-type specific binocular interactions in mouse visual thalamus

**DOI:** 10.1101/2024.07.12.603141

**Authors:** Sean P. Masterson, Govin Govindaiah, William Guido, Martha E. Bickford

## Abstract

Projections from each eye are segregated in separate domains within the dorsal lateral geniculate nucleus (dLGN). Yet, *in vivo* studies indicate that the activity of single dLGN neurons can be influenced by visual stimuli presented to either eye. In this study we explored whether intrinsic circuits mediate binocular interactions in the mouse dLGN. We employed dual color optogenetics *in vitro* to selectively activate input from each eye and recorded synaptic responses in thalamocortical (relay) cells as well as inhibitory interneurons, which have extensive dendritic arbors that are not confined to eye specific domains. While most relay cells received monocular retinal input, most interneurons received binocular retinal input; consequently, the majority of dLGN relay cells received binocular retinogeniculate-evoked inhibition. Moreover, in recordings from adjacent pairs of relay cells and interneurons, the most common relationship observed was binocular excitation of interneurons paired with binocular inhibition of adjacent relay cells. Finally, we found that dLGN interneurons are interconnected, displaying both monocular and binocular inhibition in response to retinal activation. In sum, our results indicate that geniculate interneurons provide one of the first locations where signals from the two eyes can be compared, integrated, and adjusted before being transmitted to cortex, shedding new light on the role of the thalamus in binocular vision.

**Highlights:**

- In vitro dual color optogenetics examined convergence of eye-specific retinal inputs to thalamocortical (relay) cells and interneurons in the dLGN
- The majority of relay cells receive monocular excitatory retinogeniculate input while the majority of interneurons receive binocular input
- Binocular relay cells are located in and around the ipsilateral patch whereas binocular interneurons are distributed throughout the dLGN
- The majority of relay cells receive binocular retinogeniculate-evoked inhibition
- dLGN interneurons are interconnected, receiving both monocular and binocular retinogeniculate-evoked inhibition

## Introduction

A fundamental feature of the mammalian retinogeniculate pathway is the separation of eye-specific retinal afferents in the dorsal lateral geniculate nucleus (dLGN). In carnivores and primates these inputs are arranged into retinotopically aligned, well segregated, eye-specific laminae ^1–4^. Although most rodents lack easily discernible layers within the dLGN, retinal projections are still organized into non-overlapping eye-specific domains that can be readily distinguished by anterograde labeling of retinal ganglion cell (RGC) afferent terminal fields ^5,6^. Such eye specific arrangements explain why dLGN neurons are reported to have receptive fields that are monocular, with excitatory activity driven by one eye or the other. Nonetheless, several studies across a variety of mammalian species reveal that many dLGN neurons receive some form of binocular input ^7–13^.

Studies in mice have greatly advanced our understanding of the origin and nature of binocular interactions in the dLGN. Anatomical evidence indicates that geniculocortical (relay) neurons have dendritic fields that readily cross the borders of eye specific domains ^14^. Moreover, transynaptic rabies tracing reveals that many relay cells receive direct monosynaptic input from both eyes ^15^. Indeed, some reports even suggest that almost all dLGN neurons residing in the ipsilateral eye domain receive excitatory input from the contralateral eye ^13,16^. While binocular interactions in the mouse dLGN may be more prevalent than in other species, the degree of retinogeniculate convergence appears to be highly conserved, with the excitatory input from one eye strongly dominating input from the other (nondominant) eye ^17^.

Many studies also reveal that nondominant eye input can have an inhibitory influence on dLGN activity. For example, *in vivo* recordings reveal that visual stimuli presented to the nondominant eye can suppress the visual response to the dominant eye ^7–10^. The circuitry underlying such binocular inhibitory interactions has been difficult to delineate, but both *in vitro* and *in vivo* studies suggest they are mediated by di-synaptic feedforward circuits involving inhibitory intrinsic interneurons and/or the thalamic reticular nucleus ^18–20^. In mice, intrinsic interneurons are dispersed throughout the dLGN ^21–24^, have extensive dendritic branching patterns that are not confined to eye specific domains ^21,25^ and receive a high level of retinal convergence, with some pooling input from as many as a dozen retinal ganglion cells ^25^.

While these features make interneurons excellent candidates for promoting binocular inhibitory interactions throughout the dLGN, we still know little about the prevalence, strength, and sign of their binocular responsiveness. To explore these unanswered questions, we made use of a transgenic mouse line in which green fluorescent protein (GFP) is expressed within dLGN interneurons (GAD67-GFP), making it possible to visualize them and conduct whole cell recordings in thalamic slices ^21,25^. By adopting a dual-color opsin technique, we were able to independently examine the light-evoked synaptic responses that originated from one eye or the other.

Our results reveal that while most relay cells receive retinal input from only one eye, most interneurons receive binocular input and, correspondingly, most relay cells are inhibited by stimulation of input from either eye. Finally, our results revealed abundant interconnections between inhibitory intrinsic interneurons. Collectively, these results indicate that although dLGN relay cells are primarily monocular in terms of their direct responses to input from either eye, their responses can readily be modified by input from either eye via inhibition and/or disinhibition. Thus, binocular rivalry and/or fusion may be initiated via intrinsic dLGN circuits.

## Results

### Dual optogenetic activation of ipsilateral and contralateral retinogeniculate terminals

To examine the degree to which geniculocortical (relay) cells and interneurons receive input from ipsilateral or contralateral retinogeniculate terminals, we carried out *in vitro* experiments using a dual optogenetic activation approach ^26^ in dLGN slices obtained from GAD67-GFP mice. We injected a virus in one eye to induce the expression of the red-shifted opsin Chrimson (ChRred) and a virus in the other eye to induce the expression of the blue-light activated opsins Chronos or Channelrhodopsin (ChRblue; **Figure 1A**). Two weeks later, slices of the dLGN were prepared (**Figure 1B**) and whole cell recordings were targeted to interneurons (which contain GFP in the GAD67-GFP line; **Figure 1C, i**) or relay cells (which do not contain GFP; **Figure 1C, r**). The location of the patch pipette was photographed at the conclusion of each recording, and biocytin was included in the internal solution so that the neuron identity (interneuron, **Figure 1D**, or relay cells, **Figure 1E**) could be confirmed and the location of all recorded neurons (n = 123 relay cells, n = 110 interneurons) could be plotted on a template of the dLGN. Recorded neurons were distributed across the dLGN but were concentrated in the binocular zones (dorsomedial half, indicated by the dotted line in **Figure 1F,G).**

**Figure 1.**
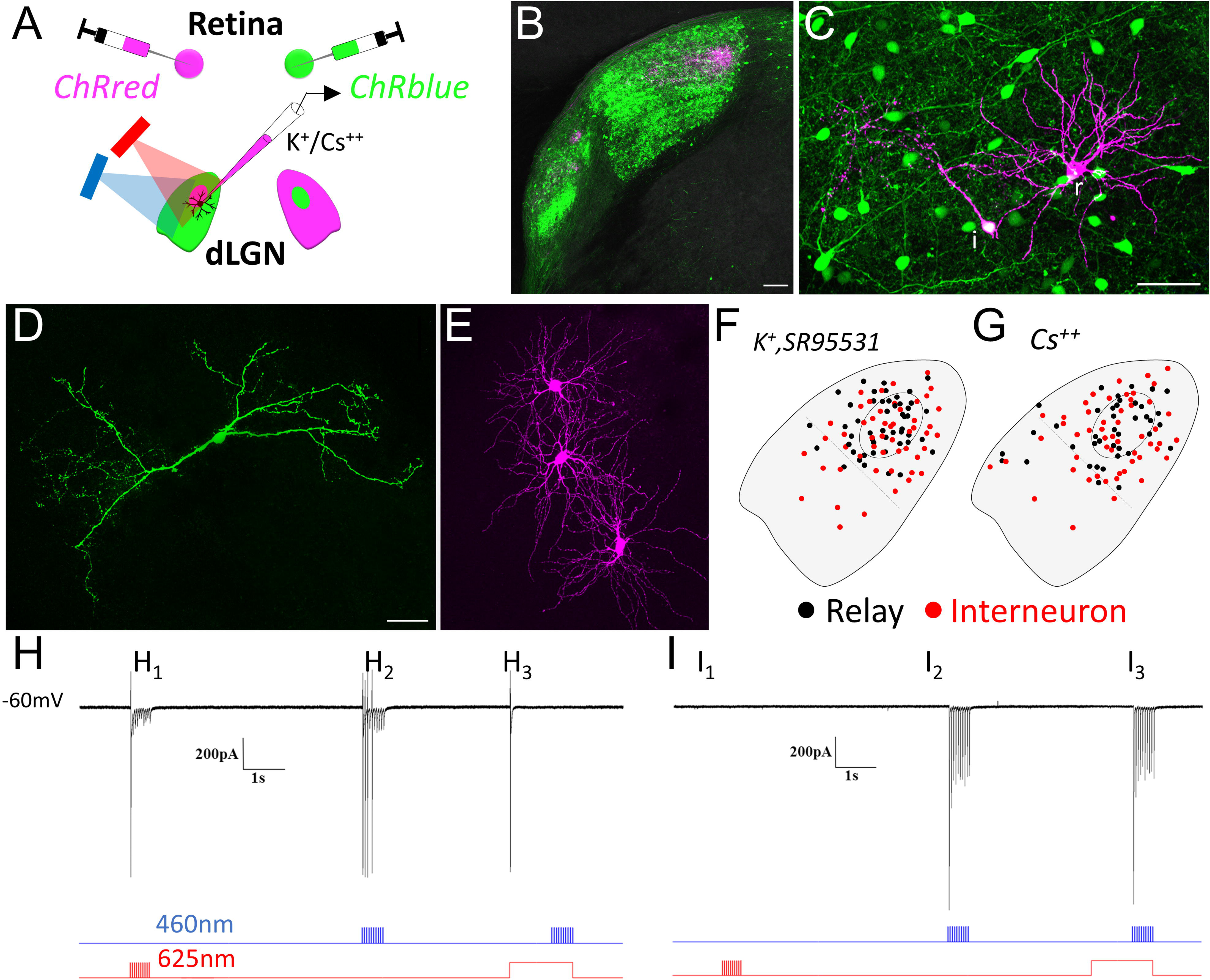
Experimental protocol. **A)** Schematic showing the experimental design and methods. In GAD67-GFP mice, a virus (pAAV9-Syn-Chrimson-tdT) was injected in one eye to induce the expression of the red-shifted opsin Chrimson (ChRred), a second virus (pAAV9-Syn-Chronos-GFP or AAV9-hSyn-hCHR2(H134R)-EYFP) was injected in the other eye to induce the expression of the blue-light activated opsins Chronos or Channelrhodopsin (ChRblue), and *in vitro* whole cell recordings were obtained from acutely prepared coronal thalamic slices. Retinal terminals were photo-stimulated (using 460nm, and/or 625nm light pulses) and light evoked synaptic responses were obtained from relay neurons and interneurons using pipettes containing K^+^ or Cs^++^ based internal solutions. **B)** Example of a coronal slice showing the viral labeling pattern of retinal terminals. Viral injections led to labeling of contralateral (green) and ipsilateral (magenta) retinal terminals in the dLGN and vLGN. **C)** Confocal image showing GFP-expressing interneurons along with a biocytin filled interneuron and relay cell. Recordings were targeted to interneurons (which contain GFP in the GAD67-GFP line; **i**) or relay cells (which do not contain GFP; **r**) and biocytin was included in the internal solution to fill the dendritic arbors (magenta). **D-E)** Confocal images show a biocytin filled interneuron (D) and 3 biocytin filled relay cells (E). **F-G)** Templates of the dLGN showing the approximate location of all recorded relay neurons (black dots) and interneurons (red dots). Each dot represents a single neuron. F) Locations of recorded relay cells (n= 49) and interneurons (n =52) recorded using K^+^ based internal solution and the GABA_A_ antagonist SR95531 in the bath. G) Locations of recorded relay cells (n=74) and interneurons (n=58) recorded using Cs^++^ based internal solutions and no GABA antagonist in the bath. **H)** Postsynaptic currents recorded in a relay cell that responded only to photoactivation of retinogeniculate terminals expressing ChRred. Traces below the response illustrate the timing, duration, and temporal frequency of blue and red light photoactivation. The ChRred was activated by 20Hz red light pulses (**H_1_**), 20Hz blue light pulses (**H_2_**) and to the initiation of a 1.5 second red light pulse which subsequently occluded responses to paired 20Hz blue light pulses (**H_3_**). **I)** Postsynaptic currents recorded in a relay cell that responded only to photoactivation retinogeniculate terminals expressing ChRblue. The ChRblue was not activated by 20Hz red light pulses (**I_1_**) but was activated by 20Hz blue light pulses (**I_2_**) even when paired with a 1.5 second red light pulse (**I_3_**). Scale bars: B = 100 um, C= 50 um, D = 50 um and also applies to E.

To activate ipsilateral or contralateral ChRred-expressing retinogeniculate terminals, 20 Hz red light pulses (10ms) were directed through the microscope objective (**Figure 1H**_1_). Since ChRred-expressing terminals are also activated by blue light ^26^ (**Figure 1H_2_**), to activate ipsilateral or contralateral ChRblue-expressing retinogeniculate terminals, a 1.5 second red light pulse was paired with 20Hz blue light pulses (1ms duration, 10 pulses). The long red-light pulse served to occlude any responses evoked from activation of the ChRred-expressing terminals; after an initial release of neurotransmitter from the ChRred-expressing terminals, the continuous light stimulation resulted in no further neurotransmitter release ^27^ (**Figure 1H_3_**). However, the red light had no effect on the ChRblue-expressing retinogeniculate terminals so that blue light pulses paired with red light could be used to exclusively activate the ChRblue-expressing terminals (**Figure 1I_3_**).

#### Excitatory retinogeniculate responses in relay cells and interneurons

Using our dual optogenetic activation technique, we first compared the direct innervation of relay cells and interneurons by ipsilateral and/or contralateral retinogeniculate terminals. For these experiments, the GABA receptor antagonist SR95531 (20 μM) was included in the bath to block interneuron synaptic activity and neurons were held at -60mV in voltage and/or current clamp mode using pipettes filled with a potassium-based internal solution. Under these conditions, monocular (**Figure 2A**) and binocular (**Figure 2B**) excitatory responses were recorded in both relay cells and interneurons. As previously described ^17^, input from one eye dominates the excitatory retinogeniculate responses of most relay cells, with 29/49 (59%) receiving monocular input. The remaining 20 (41%), received excitatory input from both eyes (**Figure 2C**), but the amplitudes of excitatory postsynaptic currents (EPSCs) were larger in response to activation of input from one eye (dominant eye) relative to the other (non-dominant) eye (**Figure 2D,H**, paired t-test, t=4.047, df=19, p<0.0007). On average dominant eye responses were 4-fold higher than nondominant eye responses (**Figure 2J**).

**Figure 2.**
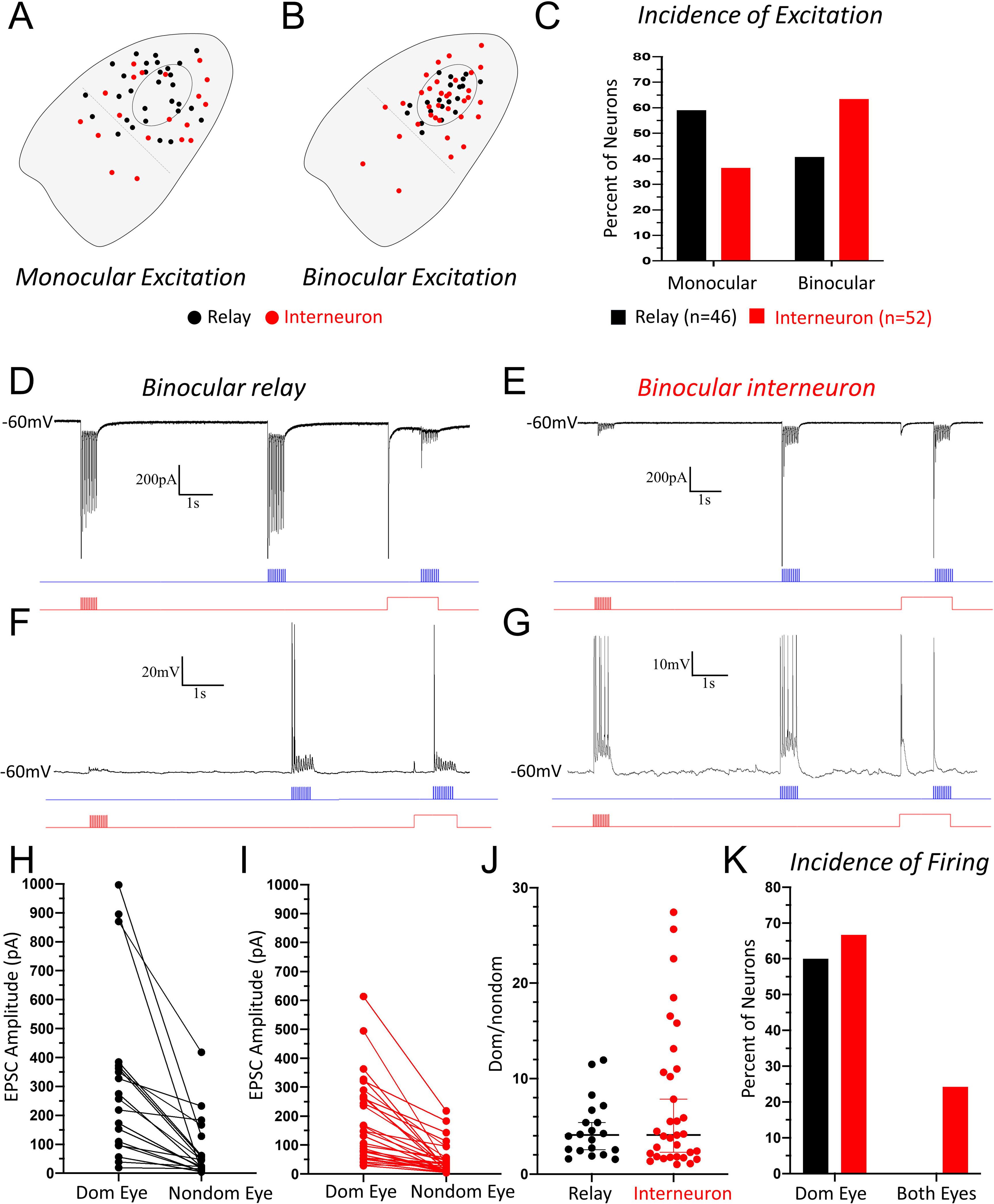
Cell-type specific excitatory postsynaptic responses evoked by photoactivation of ipsilateral and contralateral retinogeniculate terminals. **A)** Locations of recorded relay cells (black dots) and interneurons (red dots) that responded to activation of retinogeniculate terminals originating from one eye only. Monocular relay cells and interneurons are dispersed across the dLGN. **B)** Locations of recorded relay cells and interneurons that responded to activation of retinogeniculate terminals originating from both eyes. Binocular relay cells are located in and around the ipsilateral patch (small oval) while binocular interneurons were more widely dispersed. **C)** Histogram illustrating the percentages of relay cells (black bars) and interneurons (red bars) with a monocular or binocular response. Most relay cells are monocular, while most interneurons are binocular. **D)** Excitatory postsynaptic currents (EPSCs) recorded in a relay cell that responded to photoactivation of input from both eyes. **E)** EPSCs recorded in an interneuron that responded to photoactivation of input from both eyes. **F)** Excitatory postsynaptic potentials (EPSPs) recorded in a binocular relay cell; action potentials were initiated in response to input from one eye only. **G)** Excitatory postsynaptic potentials (EPSPs) recorded in a binocular interneuron; action potentials were initiated in response to input from both eyes. **H)** Plot of the EPSC amplitudes recorded in binocular relay cells. EPSCs from one eye are much stronger (dominant) than the other. Connected dots depict a single neuron’s dominant and nondominant eye response. **I)** Plot of the EPSC amplitudes recorded in binocular interneurons. Similar to relay cells, EPSCs from one eye dominate. **J)** Ratios of dominant/non-dominant EPSC amplitudes recorded in binocular relay cells and interneurons. Ratios for relay cells and interneurons were not significantly different from each other. **K)** Plot showing the incidence of monocular and binocular responses leading to spike firing. The probability of retinogeniculate inputs activating action potentials was greater in interneurons.

Interestingly, the incidence of binocular responses was higher among interneurons, with 33/52 (63%) responding to input from both eyes (**Figure 2C**). Moreover, while binocular relay cells were located in and around the ipsilateral patch of retinogeniculate terminals, binocular interneurons were more widely distributed across the dLGN (**Figure 2B**). Nonetheless, similar to relay cells, the binocular responses of interneurons were dominated by one eye (**Figure 2E, I;** paired t-test, t=6.051, df=32; p<0.0001). In fact, there was no difference in the ratio of dominant/non-dominant eye responses in relay cells versus interneurons (**Figure 2J**, Mann-Whitney U-test, U=313.5, p=0.7675).

However, retinogeniculate terminal photoactivation was more likely to initiate action potentials in interneurons (**Figure 2G**) than in relay cells (**Figure 2F**). During current clamp recordings, photostimulation of retinogeniculate terminals initiated action potentials in 91% of interneurons versus 60% of relay cells. Moreover, spikes were initiated in relay cells only in response to dominate eye input, while 24% of interneurons fired in response to activation of input from the dominate or non-dominant eyes (**Figure 2K**). A likely explanation for this difference in the incidence of firing is that the average input resistance of interneurons (697±175 mΩ, n=10) is more than double that of relay cells (331±107 mΩ, n=10; see also ^28^).

#### Retinogeniculate evoked inhibitory responses in relay cells and interneurons

We next examined the inhibitory responses in relay cells (n = 74) and interneurons (n = 58) evoked by photoactivation of ipsilateral or contralateral retinogeniculate terminals (**Figure 3A**). For these experiments, we again made use of the dual opsin approach, but did not add GABA receptor antagonists to the bath.

**Figure 3.**
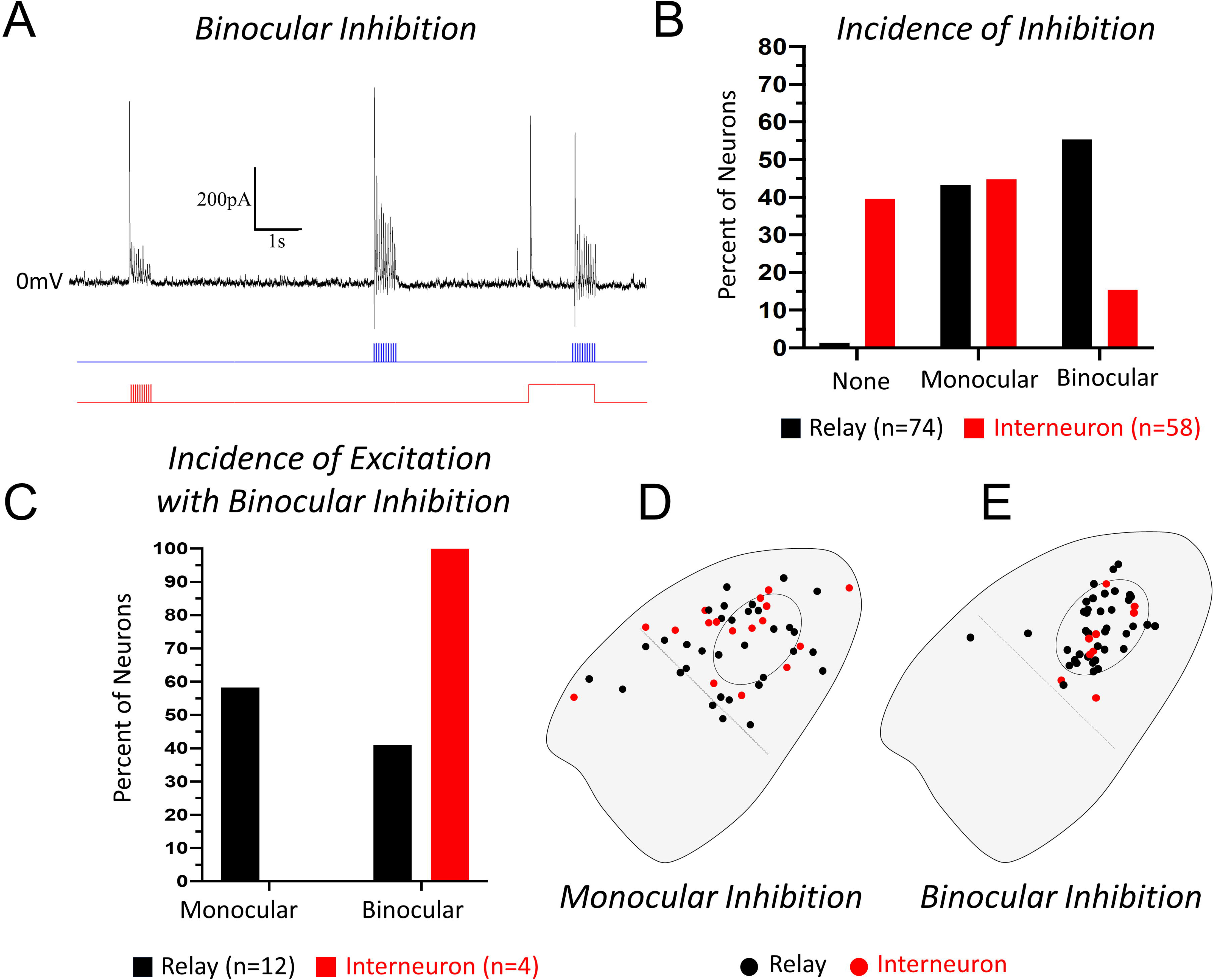
Cell-type specific inhibitory responses evoked by photoactivation of ipsilateral and contralateral retinogeniculate terminals. **A)** Inhibitory postsynaptic currents (IPSCs) recorded in a relay cell in response to photoactivation of input from either eye. **B)** Histogram illustrating the percentage of relay cells (black bars) and interneurons (red bars) with monocular, binocular, or no inhibitory responses following activation of retinogeniculate terminals. **C)** Histogram depicting the direct (monosynaptic) excitatory retinogeniculate responses recorded in relay cells and interneurons that received binocular inhibition (i.e., disynaptic IPSCs following activation of retinogeniculate input from either eye). **D)** Locations of recorded relay cells (black dots) and interneurons (red dots) that responded with IPSCs to activation of retinogeniculate terminals originating from one eye only. Relay cells and interneurons with monocular IPSCs are dispersed across the dLGN. **E)** Locations of recorded relay cells and interneurons that responded with IPSCs to activation of retinogeniculate terminals originating from both eyes. Binocular relay and interneurons are primarily located in and around the ipsilateral patch.

Additionally, pipettes were filled with a cesium-based internal solution and neurons were recorded in voltage clamp mode at holding potentials of at -60mV and 0mV. This approach allowed us to identify both excitatory and/or inhibitory currents evoked by photostimulation of retinal terminals arising from either eye.

In these experiments, nearly all relay cells (73 of 74, 99%) received some form of retinogeniculate-evoked inhibition. Reflecting the fact that most interneurons receive direct binocular innervation (**Figure 2C**), the majority (41 of 74, 55%) of relay cells exhibited inhibitory responses following activation of input originating from either eye (**Figure 3B**). Of the 41 relay cells that received binocular inhibition, only 15 (37%) were directly innervated by both ipsilateral and contralateral retinogeniculate terminals (**Figure 3C**). Moreover, 6 relay cells that received monocular inhibition did not exhibit direct excitatory responses to photoactivation of retinogeniculate terminals in the slice. Thus, despite the fact that relay cells responses are dominated by direct excitatory input from one eye, interneuron connections allow for extensive binocular inhibitory interactions within the dLGN.

The majority of interneurons (35/58, 60%) also responded with inhibitory currents following activation of retinogeniculate input, indicating that interneurons are interconnected with each other. Most of these interneurons received monocular inhibition (26, 45%; **Figure 3B**), but 9 (15%) responded with inhibitory currents following photoactivation of input from either eye. However, unlike relay cells, all of the interneurons that received binocular inhibition were also directly innervated by both ipsilateral and contralateral retinogeniculate terminals (**Figure 3C**). Furthermore, while relay cells and interneurons that received monocular inhibition were located throughout the dLGN (**Figure 2D**), those that received binocular inhibition were primarily located in and around the ipsilateral patch of retinogeniculate terminals (**Figure 2E**). This suggests that while binocular interneurons are located throughout the dLGN, their inhibitory output is spatially related to the distribution of retinogeniculate terminals that originate from either eye. In other words, the dendritic terminals of interneurons likely release GABA from sites where they receive retinal innervation.

#### Recordings from adjacent relay cell and interneuron pairs

Most of the experiments described above (using cesium-based electrodes and no GABA antagonists in the bath) were carried out by sequentially recording from adjacent relay cell and interneuron pairs. This enabled the direct comparison of the retinogeniculate responses of closely spaced neurons (i.e., in which the somata of both neurons were surrounded by the same sets of ChRred- and ChRblue-expressing retinogeniculate arbors). Given the more extensive dendritic arbors of interneurons, the dendrites of adjacent relay cells and interneurons overlapped to some degree, but often occupied distinct regions of the dLGN (**Figure 4A-C).**

**Figure 4.**
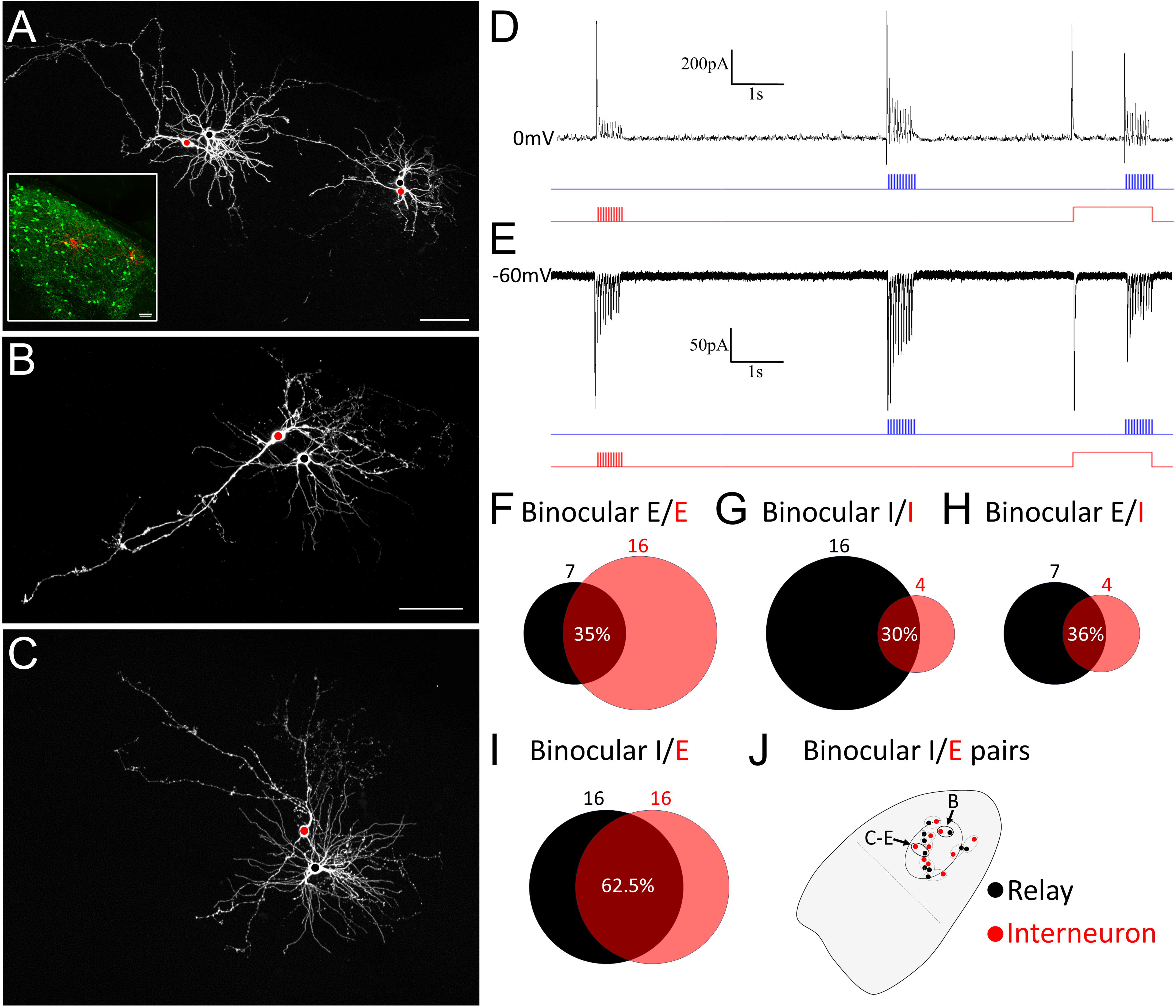
Excitatory and inhibitory responses evoked by photoactivation of ipsilateral and contralateral retinogeniculate terminals recorded in adjacent relay cells and interneurons. **A-C)** Examples of closely spaced biocytin-filled relay cells (black dot on soma) and interneurons (red dot on soma). The inset in panel A illustrates the location the 2 pairs (red) in relation to GFP-labeled interneurons in the dLGN (green). Each pair exhibits overlapping and nonoverlapping dendritic domains. **D)** Binocular IPSCs recorded in the relay cell illustrated in panel C. **E)** Binocular EPSCs recorded in the adjacent interneuron illustrated in panel C. F-I) Venn diagrams summarize the responses of relay cell/interneuron pairs that exhibited either binocular excitation (black E relay cells, red E interneurons) or inhibition (black I relay cells, red I interneurons). Black circles and numbers represent the number of relay cell responses, red circles and numbers represent the number of interneurons responses, and the overlap indicates the percentage of relay cells/interneuron pairs with identical responses (binocular excitation **F**, binocular inhibition **G**) or opposite responses (relay binocular excitation and interneuron binocular inhibition **H**, or relay cell binocular inhibition and interneuron binocular excitation, **I**). **J)** Distribution of relay cells that received binocular inhibition and adjacent interneurons that received binocular excitation. Ovals indicate relay cells/interneuron pairs; black ovals indicate pairs illustrated in panel B and in panels C-E. Scale in A = 50 um (inset 100 um). Scale in B = 50 um and also applies to C.

We recorded a total of 39 pairs of relay cells and interneurons that were closely spaced and found that they rarely displayed similar responses (**Figure 4D,E**). Seven relay cells and 16 interneurons received direct binocular retinogeniculate innervation but only 4 of these were adjacent pairs (8 of 23 cells or 35%, schematically illustrated in **Figure 4F**). Shared binocular inhibition was also rare; 16 relay cells and 4 interneurons received binocular inhibition and only 3 of these were adjacent pairs (6 of 20 cells or 30%, **Figure 4G**). The most common relationship was complementary interneuron excitation (**Figure 4E**) paired with relay cell inhibition (**Figure 4D**). For the 16 interneurons that received binocular retinogeniculate input, 10 of the adjacent relay cells received binocular inhibition (20 of 32 cells or 62.5%, **Figure 4I**). The inverse relationship was far less common; for the 7 relay cells that received binocular excitation, only 2 of the adjacent interneurons received binocular inhibition (4 of 11 cells or 36%; **Figure 4H**). Finally, the adjacent pairs that exhibited complementary binocular interneuron excitation and binocular relay cell inhibition were all located in and around the ipsilateral patch of retinogeniculate terminals (**Figure 4J**), again suggesting that interneuron output is spatially related to the retinogeniculate terminals that innervate them.

## Discussion

Five critical dLGN circuit interactions were revealed by our experiments: 1) both relay cells and interneurons receive direct binocular excitatory input from the retina, but this feature is more prevalent among interneurons, and retinogeniculate terminal photoactivation is more likely to initiate action potentials in interneurons; 2) binocular relay cells are located in and around the ipsilateral patch whereas binocular interneurons are located throughout the dLGN; 3) both cell types receive binocular retinogeniculate evoked inhibition but it is far more prevalent among relay cells than interneurons; 4) in recordings from adjacent neurons, the most common relationship observed was binocular excitation of interneurons paired with binocular inhibition of adjacent relay cells, and 5) dLGN interneurons are interconnected, displaying both monocular and binocular inhibition in response to retinal activation. Collectively, these results indicate that although the dLGN relay cells are considered monocular in terms of their direct responses to input from either eye, their responses can readily be modified by input from either eye via inhibition from interneurons and/or disinhibition brought about by interneuron-to-interneuron connections.

### Anatomical connections underlying the observed physiological responses

The physiological responses recorded in our study corroborate the known anatomical connections within the dLGN. Whyland et al., ^29^ found no significant differences in the sizes of contralateral and ipsilateral retinogeniculate terminals in the mouse and no difference in the proportion of their synaptic contacts on interneurons (ipsilateral 12%, contralateral 13%) and relay cells (ipsilateral 88%, contralateral 87%). Thus, although ipsilateral and contralateral retinogeniculate terminals originate from different subsets of retinal ganglion cells ^30^, they do not differentially innervate relay cells versus interneurons. However, since interneurons make up only 6% of neurons in the mouse dLGN ^24^, each interneuron is likely innervated by at least twice the number of retinal terminals that innervate each relay cell (also supported by available connectomic data; ^31–33)^. Moreover, relay cells primarily receive retinal input on their proximal dendrites, restricted to regions of approximately 100 um^3^ ^32,33^, while interneurons receive retinal input across their full dendritic arbors, spanning regions of up to 500 um^3^ ^21,31^. Thus, interneurons are more likely to integrate input from widely spaced retinogeniculate arbors. In addition, while there is little evidence for synaptic connections between relay cells in the mouse dLGN, recent anatomical evidence suggests that dLGN interneurons in the mouse form abundant interconnections ^31^.

Accordingly, in our study we observed that, compared to relay cells, interneurons were more likely to receive direct binocular retinogeniculate input, and most interneurons received inhibitory input from other interneurons. In addition, previous *in vitro* studies have revealed a high level of retinal convergence on interneurons ^25^.

A prominent synaptic arrangement in the dLGN is the triad. A triad is composed of a retinogeniculate terminal that synapses on interconnected relay cell dendrites and interneuron dendritic terminals (F2 profiles); i.e. an F2 profile postsynaptic to a retinal terminal synapses on a relay cell dendrite that is postsynaptic to the same retinal terminal ^34^. This synaptic arrangement likely underlies the excitatory and inhibitory currents recorded in single neurons in response to photoactivation of retinogeniculate terminals originating from the same eye. In contrast, triadic arrangements are unlikely to underlie inhibitory responses not paired with excitatory responses. These responses could be mediated via non-triadic interneuron dendritic outputs and/or interneuron axon outputs ^31^. However, given the spatial location of relay cells that received binocular inhibition, it appears that most interneuron output occurs via dendritic terminals directly innervated by retinal terminals (triadic or non-triadic). Finally, because monocular and binocular inhibitory currents were recorded in interneurons, it is possible that relay cells can be disinhibited via binocular interactions. Such a scenario is difficult to detect using our dual optogenetic activation techniques but may partially account for changes in the amplitudes of inhibitory currents with repetitive optogenetic activation of retinal terminals. In other words, strong initial inhibition of relay cells may subsequently be reduced when interneurons inhibit each other.

### Functional implications

In mice, binocular vision is essential for depth perception and acuity ^35,36^, and loss of binocular vision severely impacts fundamental behaviors such as prey capture ^30^. While cortical circuits are certainly required for creating a unified percept of visual signals originating from both eyes^37,38^, and the emergence of binocular depth perception ^35^, our study indicates that subcortical circuits, particularly those involving dLGN interneurons, may also be important components of this process. In fact, recent studies demonstrate that cortical ocular dominance plasticity is absent in adult mice lacking thalamic synaptic inhibition ^16,39^.

When one or both eyes are stimulated in awake mice, geniculate neurons exhibit dominant/non-dominate eye interactions that include both enhancement and suppression ^13^. While these interactions could be mediated by extrinsic sources (e.g. cortex, ^39^; thalamic reticular nucleus, ^40,41^; superior colliculus, ^42^; pretectum, ^43,44^; or parabigeminal nucleus, ^29,45^), our study indicates that geniculate interneurons provide one of the first locations where signals from the two eyes can be compared, integrated, and adjusted before being transmitted to cortex. Whether these interneuron-mediated binocular interactions translate to species where retinogeniculate inputs from the two eyes are more highly segregated and arranged in a laminar fashion, requires additional investigation. However, in carnivores and primates, geniculate neurons exhibit binocular suppression ^7–10^ and in both species there is evidence for interneurons with interlaminar connections ^46–48^. Although future nonrodent studies are needed, our findings shed new light on the role of the thalamus in binocular vision, providing information which may be of critical importance for the understanding and treatment of binocular vision dysfunction^49^.

## Acknowledgements

The authors thank Arkadiusz Slusarczyk and Barbara O’Steen for their assistance with histology. This work was funded by R01EY035523, R01EY031322, and R01EY12716.

## STAR Methods

### RESOURCE AVAILABILITY

#### Lead contact

Further information and requests for resources and reagents should be directed to and will be fulfilled by the lead contact, Martha Bickford.

#### Materials availability

This study did not generate new unique reagents.

#### Data availability

The datasets used and/or analyzed during the current study are available from the corresponding author on reasonable request.

### METHOD DETAILS

All breeding and experimental procedures were approved by the University of Louisville Institutional Animal Care and Use Committee. Experiments were carried out using mice, of either sex, in a line in which neurons that contain the 67KD isoform of glutamic acid decarboxylase (GAD) express green fluorescent protein (GFP; GAD67-GFP; Jax Stock No: 007677, G42 line).

#### Virus injections

To induce the expression of the opsins and fluorescent proteins in retinogeniculate axons and terminals, GAD67-GFP pups (p14-18) received intravitreal virus injections in each eye. Each pup was anesthetized with isoflurane via a small nose cone, the sclera was pierced with a sharp tipped glass pipette, and excess vitreous was drained.

Additional pipettes, filled with AAV solutions and attached to a picospritzer, were inserted into the holes made by the first pipette and volumes of approximately 1µl of AAV solution were injected into each eye. The nose cone used to administer isoflurane was then removed and once alert the pup was returned to the cage containing the dam and littermates. Intravitreal injections of pAAV9-Syn-Chrimson-tdT (Addgene) were paced in one eye to induce the expression of the red-shifted opsin Chrimson and the red fluorescent protein TdTomato in retinogeniculate terminals. In the other eye, intravitreal injections of pAAV9-Syn-Chronos-GFP or AAV9-hSyn-hCHR2(H134R)-EYFP (Addgene) were used to induce the expression of blue-light activated opsins and GFP. Initially robust expression was achieved with the first lot of pAAV9-Syn-Chronos-GFP (first 6 animals) but expression could not be achieved with subsequent lots. Therefore AAV9-hSyn-hCHR2(H134R)-EYFP was used for the final 13 animals.

#### Slice preparation and optogenetic stimulation

Two weeks following intravitreal virus injections, mice were deeply anesthetized with isoflurane and decapitated. The brain was removed from the head, chilled in cold slicing solution (in mM: 2.5 KCl, 26 NaHCO3, 2.5 KCl, 1.25 NaH2PO4, 10 MgCl2, 2 CaCl2, 234 sucrose, and 11 glucose) for 2 min, and quickly transferred into a Petri dish with room temperature slicing solution to block the brain for subsequent sectioning. Coronal slices (300 μm) through the dorsal lateral geniculate nucleus (dLGN) were cut in cold slicing solution using a vibratome (Leica VT1000 S). Slices were then transferred into a room temperature incubation solution of oxygenated (95% O2/5% CO2) artificial cerebrospinal fluid (ACSF) containing the following (in mM: 126 NaCl, 26 NaHCO3, 2.5 KCl, 1.25 NaH2PO4, 2 MgCl2, 2 CaCl2, and 10 glucose) for 30 min to 6 h. Individual slices were transferred into a recording chamber, which was maintained at 32°C by an inline heater and continuously perfused with room temperature oxygenated ACSF (2.5 ml/min, 95% O2/5% CO2). Slices were stabilized by a slice anchor or harp (Warner Instruments, Hamden, CT, United States). Neurons were visualized on upright microscopes (Olympus, BX51WI) equipped with both differential interference contrast optics and filter sets to detect fluorescence in the sections using 4X or 60X water-immersion objectives (Olympus, Center Valley, PA, United States) and a CCD camera. In GAD67-GFP mice, geniculocortical cells were identified as neurons that did not contain GFP, while interneurons were identified as cells that contained GFP.

Recording electrodes were pulled from borosilicate glass capillaries (World Precision Instruments, Sarasota, FL, United States) by using a Model P-97 puller (Sutter Instruments, Novato, CA, United States). To record excitatory postsynaptic potentials (EPSPs) or excitatory postsynaptic currents (EPSCs) in geniculate neurons, electrodes were filled with a potassium-based intracellular solution containing the following (in mM): 117 K-gluconate, 13.0 KCl, 1 MgCl2, 0.07 CaCl2, 0.1 EGTA, 10 HEPES, 2 Na2-ATP, and 0.4 Na2-GTP, with pH adjusted to 7.3 using KOH and osmolarity 290 –295 mOsm. To record inhibitory post-synaptic currents (IPSCs) as well as EPSCs in geniculate neurons, electrodes were filled with a cesium-based internal solution containing (in mM): 117 Cs-gluconate, 11 CsCl, 1 MgCl2, 1 CaCl2, 0.1 EGTA, 10 HEPES, 2 Na2-ATP, 0.4 Na2-GTP, with pH adjusted to 7.3 using CsOH and osmolarity of 290 –295 mOsm. Biocytin (0.5%) was added to both intracellular solutions to allow subsequent examination of the morphology and location of the recorded neurons.

Whole-cell recordings were concentrated in the binocular regions of the dLGN (dorsomedial half). Recordings were obtained with an Axon Instruments multiclamp 700B amplifier (Molecular Devices), and a Digidata 1440A was used to acquire electrophysiological signals. The stimulation trigger was controlled by Clampex 11.03 software (Molecular Devices). The signals were sampled at 20 kHz, and data were analyzed offline using Matlab. For current-clamp recordings, voltage signals were obtained from cells with resting potentials of −50 to −65 mV. For voltage-clamp recordings, currents were recorded at 0 mV or −60 mV.

For photoactivation of retinogeniculate terminals, light from a blue (Prizmatix UHP 460) and/or red (Prizmatix UHP 625) light-emitting diode was reflected into a 60X water-immersion objective. This produced spots of light onto the submerged slice with diameters of ∼0.3 mm (at full power, blue light 107.2 mW/mm^2^, red light 228.7 mW/mm^2^). Pulse duration and frequency were under computer control. To activate Chrimson-expressing and/or ChR2/Chronos-expressing retinogeniculate terminals the following light activation protocol was used: 500ms of 20Hz red light pulses (10ms duration), followed by 5 seconds of no light, followed by 500ms of 20Hz blue light pulses (1ms duration), followed by 3.5ms of no light, followed by 1.5 seconds of continuous red light stimulation with simultaneous 500 ms of 20Hz blue light pulses (1ms duration) during the last 500ms of the continuous red light stimulation. If light activation of retinogeniculate terminals induced spikes in the postsynaptic neurons, the light intensity was reduced to record postsynaptic responses just below the threshold for action potential activation. The location of the patch pipette was photographed at the conclusion of each recording. To block GABAergic transmission pharmacologically, in some experiments GABA receptors (GABA*_A_*) were blocked via bath application of the antagonist 2-(3-carboxypropyl)-3-amino-6-(4-methoxyphenyl)-pyridazinium bromide (SR95531, 20 μM; Tocris Bioscience, catalog #1262).

Following recording, slices were placed in a fixative solution of 4% paraformaldehyde in 0.1 M, pH 7.4 phosphate buffer (PB) for at least 24 hours. The sections were then rinsed in PB and incubated overnight in a 1:1000 dilution of streptavidin conjugated to AlexaFluor-633 (Invitrogen) in PB containing 1% Triton X-100. The following day, the sections were rinsed in PB and mounted on slides to be imaged with a confocal microscope (Olympus FV1200BX61). Confocal images of each slice were aligned with an outline of the dLGN and biocytin-filled interneurons and relay cells were plotted.

### QUANTIFICATION AND STATISTICAL ANALYSIS

Dominant and nondominant EPSC amplitudes in relay cells and interneurons were compared using paired t-tests. Dominant/nondominant EPSC ratios in relay cells and interneuron were compared using a Mann-Whitney U-test.

### KEY RESOURCES TABLE

| REAGENT or RESOURCE | SOURCE | IDENTIFIER |
| --- | --- | --- |
| Mice |  |  |
| GAD67-GFP | Jaxon Laboratories | Jax Stock # 007677, G42 line). RRID: IMSR_JAX:007677 |
| Viruses |  |  |
| pAAV9-Syn-Chronos-GFP | Addgene | Addgene Item # 59171-AA9 |
| or AAV9-hSyn-hCHR2(H134R)-EYFP | Addgene | Addgene Item # 26973-AAV9 |
| pAAV9-Syn-Chrimson-tdT | Addgene | Addgene Item # 59171-AAV9 |
| Chemicals |  |  |
| SR95531 | Tocris Bioscience | Tocris Catalogue #1262 |
| Biocytin | Sigma-Aldrich | Sigma Cat. # B4261 |
| Streptavidin-633 | Thermo Fisher Scientific | S-21375 RRID: AB_2313500 |
| Software |  |  |

## References Cited

1. Kaas, J. H., Huerta, M. F., Weber, J. T. & Harting, J. K. Patterns of retinal terminations and laminar organization of the lateral geniculate nucleus of primates. J Comp Neurol 182, 517–553 (1978).

2. Spahr, Z. R., Economides, J. R. & Horton, J. C. Retinal Input to the Primate Lateral Geniculate Nucleus Revealed by Injection of a Different Label Into Each Eye. J Neuroophthalmol 42, E596–E597 (2022).

3. Kaas, J. H., Guillery, R. W. & Allman, J. M. Some principles of organization in the dorsal lateral geniculate nucleus. Brain Behav Evol 6, 253–299 (1972).

4. Hayhow, W. R. The cytoarchitecture of the lateral geniculate body in the cat in relation to the distribution of crossed and uncrossed optic fibers. Journal of Comparative Neurology 110, 1–63 (1958).

5. Reese, B. E. ‘Hidden lamination’ in the dorsal lateral geniculate nucleus: the functional organization of this thalamic region in the rat. Brain Res 472, 119–37 (1988).

6. Kerschensteiner, D. & Guido, W. Organization of the dorsal lateral geniculate nucleus in the mouse. Vis Neurosci 34, E008 (2017).

7. Dougherty, K., et al. Binocular Suppression in the Macaque Lateral Geniculate Nucleus Reveals Early Competitive Interactions between the Eyes. eNeuro 8, (2021).

8. Guido, W., Tumosa, N. & Spear, P. D. Binocular interactions in the cat’s dorsal lateral geniculate nucleus. I. Spatial-frequency analysis of responses of X, Y, and W cells to nondominant-eye stimulation. J Neurophysiol 62, 526–543 (1989).

9. Guido, W., Tong, L., Tumosa, N., Spear, P. D. & Heidenreich, S. Binocular interactions in the cat’s dorsal lateral geniculate nucleus, II: Effects on dominant-eye spatial-frequency and contrast processing. Vis Neurosci 8, 557–566 (1992).

10. Schroeder, C. E., Tenke, C. E., Arezzo, J. C. & Vaughan, H. G. Binocularity in the lateral geniculate nucleus of the alert macaque. Brain Res 521, 303–310 (1990).

11. Sanderson, K. J., Bishop, P. O. & Darian-Smith, I. The properties of the binocular receptive fields of lateral geniculate neurons. Exp Brain Res 13, 178–207 (1971).

12. Bishop, P. O., Kozak, W., Levick, W. R. & Vakkur, G. J. The determination of the projection of the visual field on to the lateral geniculate nucleus in the cat. J Physiol 163, 503–539 (1962).

13. Howarth, M., Walmsley, L. & Brown, T. M. Binocular integration in the mouse lateral geniculate nuclei. Curr Biol 24, 1241–1247 (2014).

14. Krahe, T. E., El-Danaf, R. N., Dilger, E. K., Henderson, S. C. & Guido, W. Morphologically distinct classes of relay cells exhibit regional preferences in the dorsal lateral geniculate nucleus of the mouse. J Neurosci 31, 17437–48 (2011).

15. Rompani, S. B., et al. Different Modes of Visual Integration in the Lateral Geniculate Nucleus Revealed by Single-Cell-Initiated Transsynaptic Tracing. Neuron 93, 767–776.e6 (2017).

16. Sommeijer, J. P., et al. Thalamic inhibition regulates critical-period plasticity in visual cortex and thalamus. Nat Neurosci 20, 1716–1721 (2017).

17. Bauer, J., et al. Limited functional convergence of eye-specific inputs in the retinogeniculate pathway of the mouse. Neuron 109, 2457–2468.e12 (2021).

18. Žiburkus, J., Lo, F. S. & Guido, W. Nature of inhibitory postsynaptic activity in developing relay cells of the lateral geniculate nucleus. J Neurophysiol 90, 1063–1070 (2003).

19. Lindström, S. Synaptic organization of inhibitory pathways to principal cells in the lateral geniculate nucleus of the cat. Brain Res 234, 447–453 (1982).

20. Lindström, S. Interneurons in the lateral geniculate nucleus with monosynaptic excitation from retinal ganglion cells. Acta Physiol Scand 119, 101–103 (1983).

21. Charalambakis, N. E., Govindaiah, G., Campbell, P. W. & Guido, W. Developmental Remodeling of Thalamic Interneurons Requires Retinal Signaling. J Neurosci 39, 3856– 3866 (2019).

22. Gabbott, P. L. A., Somogyi, J., Stewart, M. G. & Hámori, J. A quantitative investigation of the neuronal composition of the rat dorsal lateral geniculate nucleus using GABA-immunocytochemistry. Neuroscience 19, 101–111 (1986).

23. Golding, B., et al. Retinal input directs the recruitment of inhibitory interneurons into thalamic visual circuits. Neuron 81, (2014).

24. Evangelio, M., García-Amado, M. & Clascá, F. Thalamocortical Projection Neuron and Interneuron Numbers in the Visual Thalamic Nuclei of the Adult C57BL/6 Mouse. Front Neuroanat 12, (2018).

25. Seabrook, T. A., Krahe, T. E., Govindaiah, G. & Guido, W. Interneurons in the mouse visual thalamus maintain a high degree of retinal convergence throughout postnatal development. Neural Dev 8, (2013).

26. Klapoetke, N. C., et al. Independent optical excitation of distinct neural populations. Nat Methods 11, 338–346 (2014).

27. Lin, J. Y. A user’s guide to channelrhodopsin variants: features, limitations and future developments. Exp Physiol 96, 19–25 (2011).

28. Jager, P., et al. Dual midbrain and forebrain origins of thalamic inhibitory interneurons. Elife 10, 1–29 (2021).

29. Whyland, K. L., Hernandez, Y., Slusarczyk, A. S., Guido, W. & Bickford, M. E. The parabigeminal nucleus is a source of “retinogeniculate replacement terminals” in mice that lack retinofugal input. Journal of Comparative Neurology 530, 3179–3192 (2022).

30. Johnson, K. P., et al. Cell-type-specific binocular vision guides predation in mice. Neuron 109, 1527–1539.e4 (2021).

31. Morgan, J. L. & Lichtman, J. W. An Individual Interneuron Participates in Many Kinds of Inhibition and Innervates Much of the Mouse Visual Thalamus. Neuron (2020) doi:10.1016/j.neuron.2020.02.001.

32. Morgan, J. L., Berger, D. R., Wetzel, A. W. & Lichtman, J. W. The Fuzzy Logic of Network Connectivity in Mouse Visual Thalamus. Cell 165, 192–206 (2016).

33. Maher, E. E., et al. 3D electron microscopy and volume-based bouton sorting reveal the selectivity of inputs onto geniculate relay cell and interneuron dendrite segments. Front Neuroanat 17, (2023).

34. Guillery, R. W. The organization of synaptic interconnections in the laminae of the dorsal lateral geniculate nucleus of the cat. Z Zellforsch Mikrosk Anat 96, 1–38 (1969).

35. Boone, H. C., et al. Natural binocular depth discrimination behavior in mice explained by visual cortical activity. Curr Biol 31, 2191–2198.e3 (2021).

36. Stephany, C. É., et al. Plasticity of binocularity and visual acuity are differentially limited by nogo receptor. J Neurosci 34, 11631–11640 (2014).

37. la Chioma, A., Bonhoeffer, T. & Hübener, M. Disparity Sensitivity and Binocular Integration in Mouse Visual Cortex Areas. J Neurosci 40, 8883–8899 (2020).

38. Fu, J., Tanabe, S. & Cang, J. Widespread and multifaceted binocular integration in the mouse primary visual cortex. Journal of Neuroscience 43, (2023).

39. Qin, Y., et al. Thalamic regulation of ocular dominance plasticity in adult visual cortex. Elife 12, (2023).

40. Campbell, X. P. W., Govindaiah, G., Masterson, S. P., Bickford, X. M. E. & Guido, W. Synaptic properties of the feedback connections from the thalamic reticular nucleus to the dorsal lateral geniculate nucleus. J Neurophysiol 124, 404–417 (2020).

41. Soto-Sánchez, C., Wang, X., Vaingankar, V., Sommer, F. T. & Hirsch, J. A. Spatial scale of receptive fields in the visual sector of the cat thalamic reticular nucleus. Nat Commun 8, (2017).

42. Bickford, M. E., Zhou, N., Krahe, T. E., Govindaiah, G. & Guido, W. Retinal and tectal “Driver-Like” inputs converge in the shell of the mouse dorsal lateral geniculate nucleus. Journal of Neuroscience 35, (2015).

43. Wang, S., Eisenback, M., Datskovskaia, A., Boyce, M. & Bickford, M. E. M. E. M. E. GABAergic pretectal terminals contact GABAergic interneurons in the cat dorsal lateral geniculate nucleus. Neurosci Lett 323, 141–5 (2002).

44. Cucchiaro, J. B. B., Bickford, M. E. E. & Sherman, S. M. M. A GABAergic projection from the pretectum to the dorsal lateral geniculate nucleus in the cat. Neuroscience 41, 213–26 (1991).

45. Sokhadze, G., Whyland, K. L., Bickford, M. E. & Guido, W. The organization of cholinergic projections in the visual thalamus of the mouse. Journal of Comparative Neurology 530, 1081–1098 (2022).

46. Sanchez-Vives, M. V., Bal, T., Kim, U., Krosigk, M. von & McCormick, D. A. Are the Interlaminar Zones of the Ferret Dorsal Lateral Geniculate Nucleus Actually Part of the Perigeniculate Nucleus? Journal of Neuroscience 16, 5923–5941 (1996).

47. Wilson, J. R. Synaptic connections of relay and local circuit neurons in the monkey’s dorsal lateral geniculate nucleus. Neurosci Lett 66, 79–84 (1986).

48. Wilson, J. R., Forestner, D. M. & Cramer, R. P. Quantitative analyses of synaptic contacts of interneurons in the dorsal lateral geniculate nucleus of the squirrel monkey. Vis Neurosci 13, 1129–1142 (1996).

49. Peli, E. & Jung, J. H. Review: Binocular double vision in the presence of visual field loss. J Vis 24, 13 (2024).

